# A light-modulated clock mechanism in a hydrozoan jellyfish that synchronises evening gamete release

**DOI:** 10.1101/2025.05.05.651927

**Authors:** Ruka Kitsui, Noriyo Takeda, Evelyn Houliston, Ryusaku Deguchi, Tsuyoshi Momose

## Abstract

For marine species that reproduce by external fertilisation, spawning is precisely coordinated within a local cohort to maximise the chances of producing offspring. Gamete release is often synchronised with respect to the daily light changes at sunrise and sunset. In the hydrozoan jellyfish *Clytia hemisphaerica*, morning spawning occurs when oocyte maturation and gamete release are induced by maturation-inducing hormone (MIH) neuropeptides released from opsin-expressing cells in the gonad directly upon light stimulus. Here, we characterise the distinct spawning cycle of a previously undescribed species *Clytia* sp. IZ-D identified on the Pacific coast of Japan, which releases gametes in the evening. *Clytia* sp. IZ-D jellyfish spawn 14 hours after light stimuli under a 24-hour light-dark cycle and exhibit autonomous and synchronised spawning cycles with a 20-hour interval under constant light. We found that the female spawning cycle reflects the growth and acquisition of maturation competence by oocytes, such that each day a new batch of growing oocytes becomes responsive to MIH at a time that correlates with the timing of actual spawning. We propose that the synchronised evening spawning in this species is controlled by an unusual circadian timing mechanism based on the progressive development of gamete competence to MIH and modulation of the opsin-controlled MIH signalling pathway. This mechanism may provide resilience to light cycle instability due to local climate variation and ensure reproductive isolation from other *Clytia* species by alternating the gamete release timing.

## Introduction

Marine organisms rely on various environmental cues to synchronise sexual behaviour and gamete release, thereby maximising the chances of successful sexual reproduction. Precise synchronisation of spawning timing often depends on daily changes in sunlight luminosity, which serve as reliable external triggers for marine animals. For instance, dozens of coral species release gametes simultaneously within a few hours after sunset, a few nights after the full moon in late spring on the Great Barrier Reef. This occurs despite the risk of species hybridisation (Babcock et al., 1986; Harrison et al., 1984), likely using sunlight as the primary external signal for synchronising gamete release. For many marine animals, spawning control is complex, integrating multiple environmental cues, and its molecular basis remains uncharacterised. For example, annual coral gamete release is influenced by wind (Woesik, 2010), solar irradiation (Woesik et al., 2006), temperature (Babcock et al., 1986; Keith et al., 2016; Nozawa, 2012), and the lunar cycle (Harrison et al., 1984; Kaniewska et al., 2015; Shlesinger and Loya, 1985), in addition to the daylight cycle (Harrison et al., 1984; Levitan et al., 2011).

The molecular basis of light-controlled gamete release and preceding oocyte maturation is relatively well described in hydrozoan jellyfish, where daily gamete release is often directly triggered by light stimuli (Miller, 1979). In *Cytaeis uchidae* and *Clytia hemisphaerica*, exposure to light after a few hours of darkness almost immediately triggers oocyte meiotic maturation (Freeman, 1987; Freeman and Ridgway, 1988; Takeda et al., 2006). Eggs are released as meiotic divisions are completed, slightly after sperm release from male jellyfish gonads. Both processes are initiated by light-induced secretion of a maturation-inducing hormone (MIH) from neural-type cells in the gonad ectoderm. MIH comprises short neuropeptides (e.g., WPRPamide, RPRPamide) common between hydrozoan jellyfish *C. hemisphaerica* (Leptothecata) and *Cladonema pacificum* (Anthoathecata). These MIH-secreting cells also express opsin genes (Quiroga-Artigas et al., 2018). In *C. hemisphaerica*, knockout of *CheOpsin9*, which is predominantly expressed in the gonad, prevents light-triggered MIH release. Secreted MIH binds to a G-protein-coupled seven-transmembrane receptor (MIH-R) on the oocyte surface, triggering oocyte maturation via Gαs, cAMP, and cAMP-dependent protein kinase signalling (Takeda et al., 2006).

Spawning in some hydrozoan jellyfish populations can be triggered by light-to-dark transitions rather than dark-to-light, as seen in subspecies of *Cladonema pacificum* and *Spirocodon saltatrix* (Deguchi et al., 2005; Ikegami et al., 1978), or by both transitions in some *Clytia* populations (Roosen-Runge, 1962). In ‘dark-type’ female *Cladonema*, a 3–5-minute dark pulse promotes oocyte maturation, with spawning occurring after 35 minutes. These jellyfish may employ different opsins and downstream signal transduction pathways to trigger MIH release from neurons. In both light- and dark-type jellyfish, oocyte maturation is initiated by MIH release, triggered by light or dark stimuli, respectively (Takeda et al., 2018).

In addition to mechanisms that promote immediate gamete release upon light or dark stimuli, circadian clock mechanisms may also regulate spawning timing. Light-modulated circadian behaviours have been described in several cnidarians, suggesting that circadian clocks could contribute to spawning synchronisation. Both *Hydra* (Hydrozoa) polyps and the upside-down jellyfish *Cassiopea* (Scyphozoa) exhibit cyclic sleep-like state changes (Kanaya et al., 2020; Nath et al., 2017), while calcification in the reef coral *Acropora eurystoma* (Anthozoa) occurs during the day and maintains circadian oscillations under constant light (Gutner-Hoch et al., 2016). The sea anemone *Nematostella vectensis* (Anthozoa) exhibits differential locomotive behaviours between day and night, which persist under constant darkness (Oren et al., 2015). Furthermore, components of evolutionarily conserved circadian clock genes, *Clock* and *Cycle* (*BMAL1*), have been identified in anthozoans (Hoadley et al., 2011; Reitzel et al., 2010). The *Clock* orthologue in *Nematostella*, *NvClk*, is required for maintaining circadian locomotor rhythms under constant light (Aguillon et al., 2024).

Despite these findings, circadian clocks have not previously been implicated in the temporal control of oocyte maturation and gamete release in hydrozoans. Orthologues of *Clock* and *Cycle* (*BMAL1*) appear absent from the genomes of *Hydra* and *Clytia hemisphaerica* (Kanaya et al., 2019) and were likely lost in the hydrozoan common ancestor. Additionally, the transcription-translation feedback loop model underlying CLOCK-driven circadian clocks may lack the precision needed to trigger rapid events such as oocyte maturation or gamete release, which occur within minutes in hydrozoans. Moreover, hydrozoan gamete release can be directly altered by manipulating light–dark cycles, provided a minimum latent interval is respected (Takeda et al., 2006).

Here, we present the first evidence of an autonomous clock mechanism precisely controlling and synchronising the timing of oocyte maturation and gamete release in the hydrozoan jellyfish *Clytia* sp. IZ-D, a close relative of the model species *Clytia hemisphaerica*. In natural conditions, gamete release in this species occurs shortly after sunset. Through a series of experiments using intact jellyfish and isolated oocytes, we define how light separately regulates both oocyte growth and the precise timing of spawning in *Clytia* sp. IZ-D, and propose a simple model for this circadian clock-independent nightly gamete release.

## Results

Jellyfish of a new *Clytia* species (*Clytia* sp. IZ-D) were collected on Izushima Island (N:38.440501, E: 141.525718), Miyagi, along the Pacific coast of Japan (Fig. 1A) on 12 October 2023. The water temperature was 21.3°C at 2 metres depth. The average sea surface water temperature in this area in mid-October from 2014 to 2023 was 18.9°C (Fig. 1B), indicating elevated seawater temperature in 2023. Daylight duration was approximately 11.5 hours (Fig. 1C). We established a female strain (D-A1) and a male strain (D-A3) as vegetatively grown polyp colonies from planula larvae siblings, offspring of a pair of male and female jellyfish kept in a plastic dish. We reared clonal jellyfish liberated from these colonies for use in this study. In the same location, other species, notably *Clytia* sp. IZ-C (collected in July 2023, sea surface temperature 19.4°C) was also sampled. Taxonomic analysis using maximum likelihood topologies of the mitochondrial 16S rRNA gene indicated that *Clytia* sp. IZ-D is distinct from any previously barcoded *Clytia* species, including the hydrozoan model species *Clytia hemisphaerica* (Fig. 1D). However, the morphology and life cycle of *Clytia* sp. IZ-D (Fig. 1E, Suppl. Fig. 3) were indistinguishable from those of *C. hemisphaerica*. The life cycle consisted of a polyp colony with multiple feeding polyps (gastrozooids) and jellyfish budded from gonozooid polyps, which grow and mature within 10 days.

**Figure 1.**
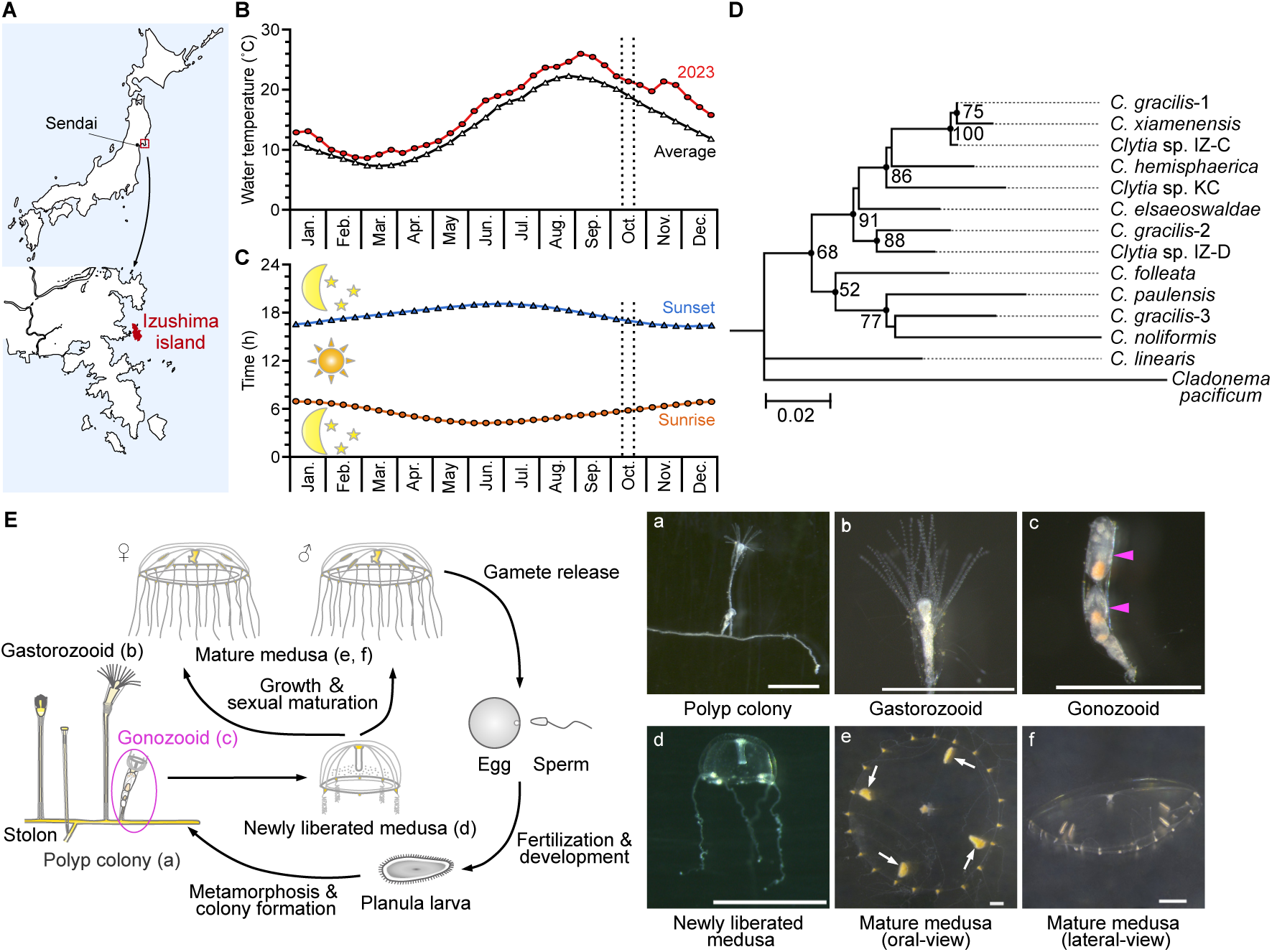
*Clytia* sp. IZ-D from Izushima island, on the Pacific coast of Japan. (A) Geographical location of the sampling. (B) Annual sea surface temperatures in 2023 (sampled year) and 10-year average, observed at Enoshima Island, 8 km southeast of Izushima. (C) Annual change of the sunrise and sunset times. The sampling period (mid-October in 2023) is indicated by vertical dotted lines in B, and C. (D) Molecular phylogeny of 16S rRNA sequence from *Clytia* species obtained by Maximum-Likelihood analysis. Bootstrap values of 1000 pseudoreplicates above 50% were shown as node-support values. (E) Life-cycle diagram (left) and images of each stage (a–f) of *Clytia* sp. IZ-D, which is morphologically very similar to the model species *Clytia hemisphaerica*. Magenta arrows in the gonozoid image (c) indicate individual medusa buds. Arrows in mature female medusa (e) point to the four gonads in the subumbrella.

*Clytia* sp. IZ-D exhibited physiological differences from *C. hemisphaerica*. Notably, this species releases gametes at night in the laboratory at the standard culture temperature of 21°C, while *C. hemisphaerica* releases eggs and sperm within 100–120 minutes after exposure to light following 3 hours of dark (Amiel and Houliston, 2009).

### Sunrise triggers *Clytia* sp IZ-D ovulation the following evening

To investigate how gamete release is controlled in *Clytia* sp. IZ-D jellyfish, we primarily focused on ovulation (egg spawning) in females using strain D-A1. Jellyfish were maintained under standard 12-hour light/12-hour dark (L12/D12) cycles at 21°C. These environmental regimes may be referred to as Zeitgebers. Under these conditions, eggs containing a clearly visible female pronucleus, indicating completion of meiosis, were released from the gonad (ovary) approximately two hours after the light-to-dark transition (Fig. 2A, B; Suppl. Video 1). We defined the gamete release time of an individual jellyfish as the moment when the first egg or sperm release was observed.

**Figure 2.**
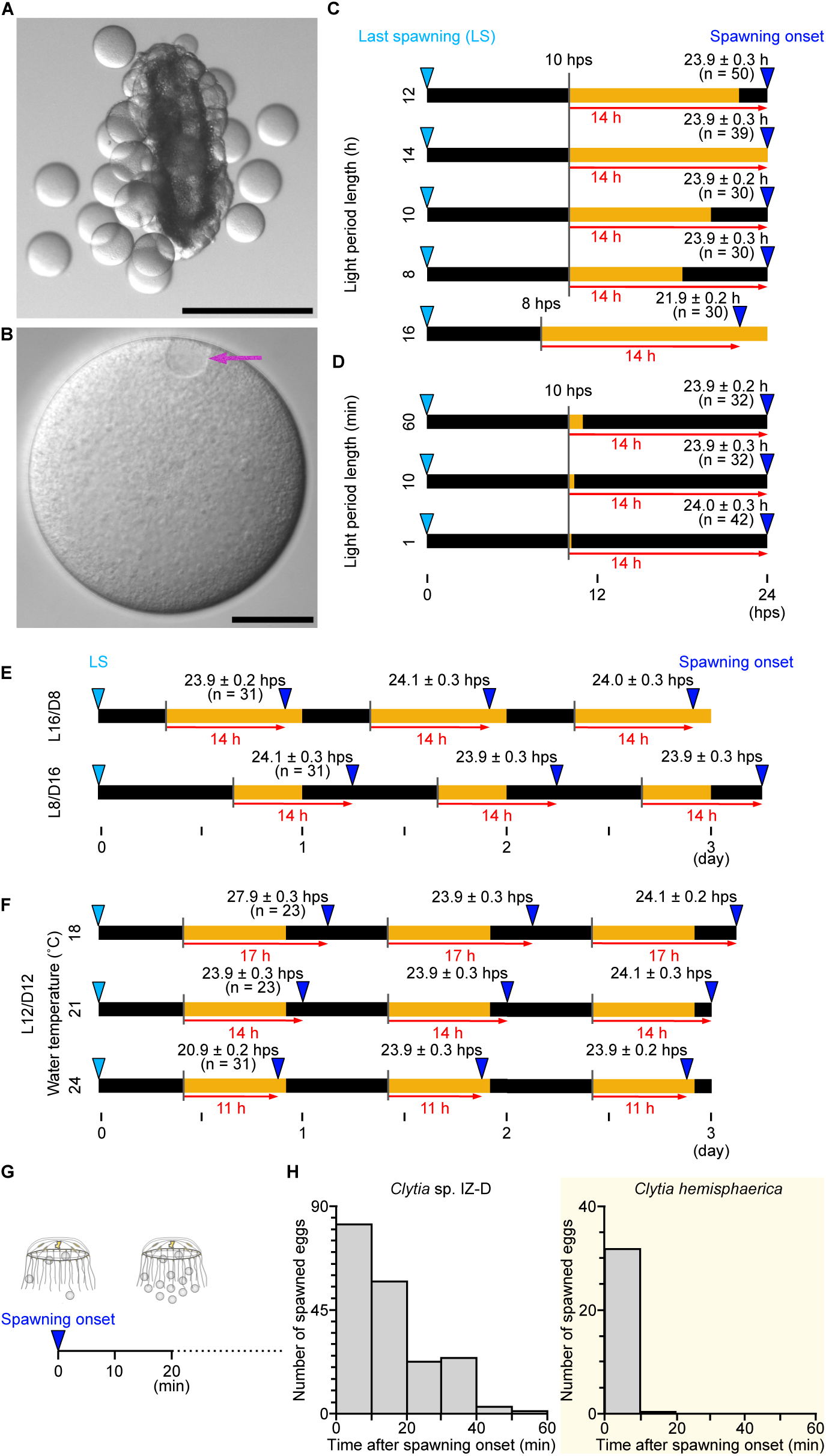
Ovulation occurs 14 hours after the light stimulus at 21°C. (A) Snapshot image of eggs being released from an isolated female gonad, which occurs with the same timing as in intact jellyfish. Scale bar: 0.5 mm (B) A mature unfertilised egg released from the gonad. Magenta arrow indicates the female pronucleus. Scale bar: 50 µm (C) The timing of spawning onset when the start or end of the light period was changed; spawning occurs 14 hours from the onset of light. (D) A short light period was sufficient to induce spawning 14 hours after light onset. (E) Daily regular spawning of jellyfish maintained under a constant Zeitgeber; spawning started about 14 hours after each light stimulus, occurring either before or after the end of the light period, depending on its length. (F) Temperature dependence of the interval between light stimuli and spawning. Jellyfish spawned regularly 17, 14 and 11 hours after the light stimulus at 18, 21 and 24°C, respectively, under an L12/D12 cycle. For all experiments shown in (C–F), jellyfish were maintained in L12/D12 at 21°C until the light cycle or temperature was changed just after the last spawning (LS: indicated with light blue triangles). Horizontal bars represent the light and dark periods, with LS as the origin of the time scale. Key intervals from the light stimuli to the spawning, likely constraining the spawning schedule, are annotated with red arrows with approximate lengths in hours; *n* =number of jellyfish used for each condition. (G) Experimental design for individual oocyte release timing measurement. The number of eggs released from a jellyfish was counted every 10 minutes following the first egg release. (H) Timing of individual oocyte release from the onset of ovulation for each jellyfish, comparing *Clytia* sp. IZ-D (N = 10) and *Clytia hemisphaerica (*N = 10).

Initially, we hypothesised that gamete release in *Clytia* sp. IZ-D was induced by the onset of darkness, similar to reports for a night-spawning subspecies of *Cladonema pacificum* (Takeda et al., 2006) and certain *Clytia* species from the North Pacific (Roosen-Runge, 1962). Contrary to our expectations, altering the onset of the dark period (dusk) did not affect the ovulation timing (Fig. 2C). However, advancing the start of the light period (dawn) by two hours (to 8 hps) led to ovulation occurring two hours earlier. In all tested cases in the Fig. 2C, ovulation occurred 14 hours after the preceding dark-to-light transition, even the light period was extremely short (as brief as one hour or one minute; Fig. 2D). These observations strongly suggest that the dark-to-light transition, which we hereafter refer as “light stimulus”, acts as a trigger for ovulation with a 14-hour delay. Such a delayed response has not been described previously. When jellyfish were subjected to Zeitgebers with different daytime lengths (8 and 16 hours of light), ovulation consistently occurred 14 hours after the light stimulus, resulting in a 24-hour cycle (Fig. 2E). Under L16/D8 cycles (16 hours of light and 8 hours of dark), ovulation occurred during the light phase (two hours before sunset). This confirms that the light stimulus, and not the light-to-dark transition, is responsible for inducing ovulation. The onset of ovulation for most jellyfish varied by only ±0.3 hours among individuals cultured under the same conditions (Fig. 2C–E). This synchronisation is notable given the 14-hour delay between the light stimulus and ovulation. To explore the precision of this delay and the factors influencing it, we examined Zeitgebers involving switches to higher (24°C) or lower (18°C) incubation temperatures (Fig. 2F) under the standard L12/D12 cycle. Ovulation occurred 11 or 17 hours after the light stimulus at 24°C and 18°C, respectively, compared to 14 hours at 21°C. Despite this temperature-dependent delay, the variance in ovulation timing within each cohort remained low (±0.2–0.3 hours). We conclude that the light stimulus, corresponding to sunrise in the natural habitat, robustly determines the timing of ovulation in the evening in a well-synchronised manner. Similarly, male jellyfish (D-A3) released sperm 13 hours after the light stimulus (Suppl. Fig. 1; Suppl. Video 2) at 21°C.

These results collectively indicate that *Clytia* sp. IZ-D possesses an unusual “delayed response” timer mechanism for synchronising evening gamete release. We further noticed that egg release from *Clytia* sp. IZ-D jellyfish continued for up to 60 minutes from its onset, compared to 10 minutes for the morning-spawning *C. hemisphaerica* (Fig. 2E, F). This difference suggests that in *Clytia* sp. IZ-D, the timing of oocyte maturation induction may be partly controlled independently among oocytes, whereas in C. *hemisphaerica*, MIH release is the dominant factor (Quiroga-Artigas et al., 2018).

### *Clytia* sp. IZ-D ovulation is regulated by a light-modulated clock

Ovulation induced directly by a light-to-dark transition is a common strategy for species that release gametes in the evening (Deguchi et al., 2005; Ikegami et al., 1978). Based on the observations presented above, we speculated that the extended delay from the light stimulus to ovulation observed in *Clytia* sp. IZ-D reflects a distinct circadian mechanism regulating gamete release, involving factors that operate autonomously within individual oocytes. To test this hypothesis, we placed female jellyfish under constant light. Interestingly, *Clytia* sp. IZ-D spawned eggs every 19 - 20 hours for at least three ovulation cycles at 21°C (Fig. 3A). The interval of ovulation was temperature-dependent, occurring every 17–19 hours and 20–22 hours at 24°C and 18°C, respectively (Fig. 3A), matching the delay of ovulation from the light stimuli under the Zeitgeber with a light-dark cycle. Across all temperature conditions, the variance in the onset of the ovulation was greater (up to ±0.9 hours) under constant light than under the Zeitgeber with a light-dark cycle, particularly during the second or third ovulation cycle. This increased variance may partly reflect the accumulation of variation over successive ovulation cycles. However, we observed that egg release from a single jellyfish continued longer, up to 80 minutes, under constant light (Fig. 3B). This supports the speculation that the autonomous ovulation cycle is controlled at the level of individual oocytes. Altogether, we conclude that *Clytia* sp. IZ-D has autonomous quasi-daily gamete release cycles operating at around 20 hrs per cycle under constant light. A reliable daily dark-light cue will modulate ovulation timing to a 24-hour cycle, synchronising gamete release among jellyfish under the same environmental conditions and among oocytes within an individual jellyfish.

**Figure 3.**
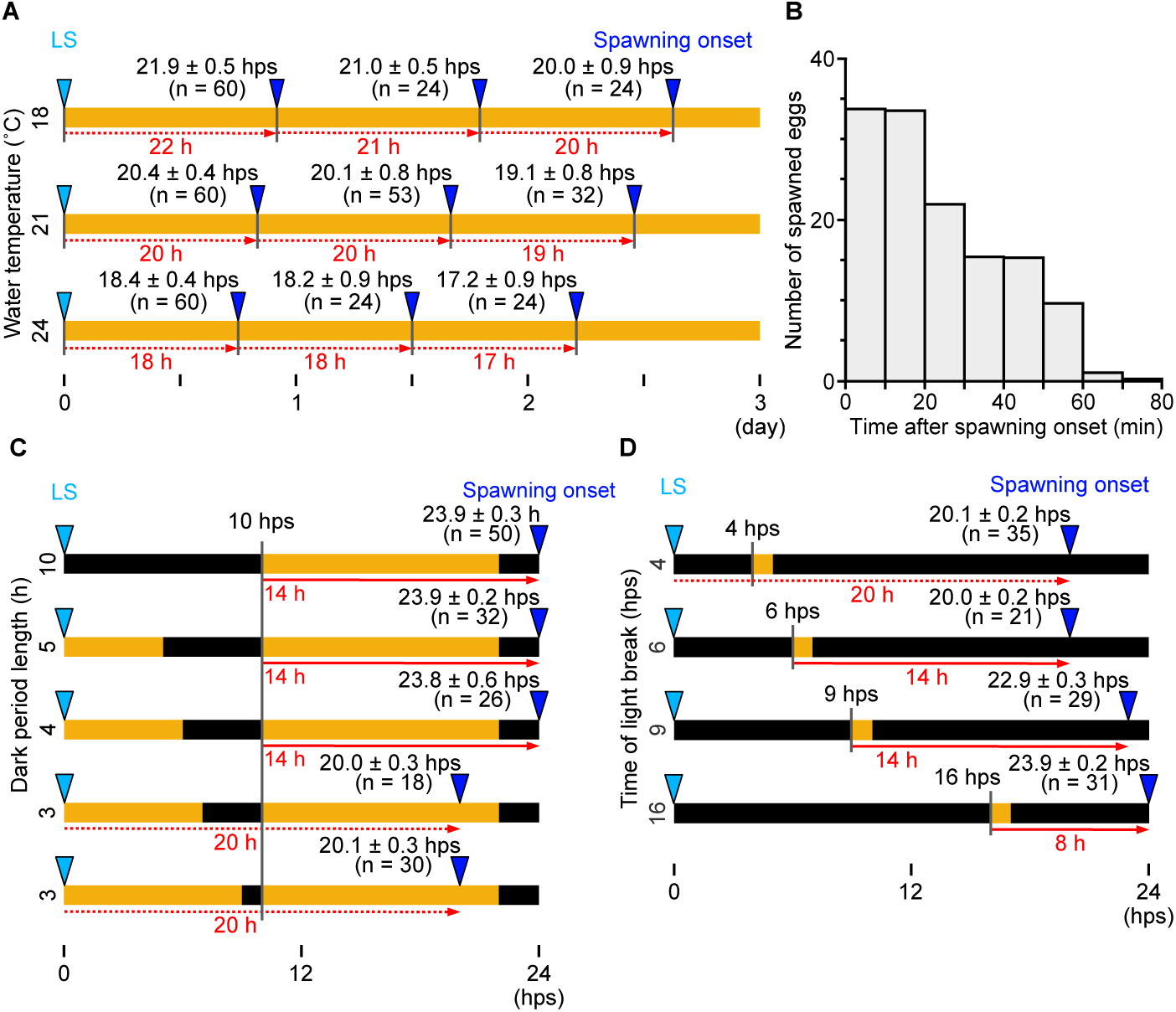
Autonomous synchronous ovulation cycles under constant light in *Clytia* sp. IZ-D. Spawning behaviour was monitored following different light-dark regimes, represented graphically as in Figure 2. (A) Cycles of synchronous egg spawning over three days under constant light. (B) Individual egg release from jellyfish (N = 10) under constant light, measured from the onset of spawning for each jellyfish. (C) Demonstration of a 4-hour minimum dark period required before the dark-to-light transition, so that it modulates the spawning as a light stimulus. (D) Influence of light stimulus timing on spawning. Spawning occurred at 20 hps when the light stimulus was applied 6 hps or earlier, and not later than 24 hps when applied 10 hps or later. Egg spawning is thus constrained to a window between 20 and 24 hps. In (A), (C) and (D), jellyfish were maintained in the standard Zeitgeber of L12/D12 at 21°C and shifted to the experimental conditions after the last spawning (LS). Red dotted arrows and numbers indicate the interval between each spawning. Red arrows and numbers represent the key approximate intervals between the light stimulus and the spawning, when the spawning cycle differs from the autonomous spawning cycle (i.e. 20 hps at 21°C).

The observation that ovulation induced by light stimuli occurs later than for jellyfish placed under constant light, i.e. without light stimuli, seemed curious if the dark light stimulus were indeed a positive trigger for ovulation. To examine how the light stimulus modulates the autonomous ovulation cycle, we altered the duration of the dark period in the standard Zeitgeber while maintaining the timing of the dark-to-light transition after the previous ovulation (i.e., 10 hps; Fig. 3C). As observed previously, the dark-to-light shift induced (delayed) ovulation after 14 hours when the dark period was four hours or longer (Fig. 3C). In contrast, ovulation occurred at 20 hps when the dark period was three hours or shorter, matching the timing observed under constant light. This indicates that a four-hour dark period is necessary for the dark-to-light transition to act as an effective light stimulus. Similarly, minimum dark periods are required for jellyfish that spawn eggs immediately after the light stimulus, including *C. hemisphaerica* (Amiel and Houliston, 2009; Takeda et al., 2006), probably to allow recovery or re-sensitisation of opsins to light.

We then addressed how the light stimulus interacts with the 20-hour autonomous ovulation cycle. For instance, the action of the light stimulus in *Clytia* sp. IZ-D could be explained either by a delaying effect on the autonomous cycle or by resetting or overriding the cycle to initiate a new 14-hour timer. To address this, we substantially altered the timing of the light stimulus, following a preceding dark period of four hours or more (Fig. 3D). A light stimulus earlier than 6 hps induced ovulation at 20 hps, equivalent to the autonomous ovulation cycle under constant light. Light stimuli between 6 and 10 hps induced ovulation 14 hours later, as characterised under the standard light cycle (Fig. 2A). A third situation was observed when the light stimulus was applied 10 hps or later, with ovulation occurring at 24 hps, less than 14 hours after the light stimulus (Fig.3D). This last observation is inconsistent with the model that the light stimulus overrides the autonomous ovulation cycle and starts a 14-hour timer. The effect of the light stimulus is thus better described as delaying the autonomous 20-hour ovulation cycle, acting within a light-sensitive window between 6 to 10 hps.

### Asynchronous ovulation under constant dark suggests a permissive role of light to maintain the circadian clock

The results presented above collectively indicate that daily light cues modulate the autonomous quasi-circadian gamete release cycle. However, light also appears to play a permissive role in maintaining the cycle, such that jellyfish release eggs asynchronously when kept under constant dark conditions. We removed jellyfish from constant-dark conditions after either 20, 24, or 28 hps and counted the number of eggs already released, or released within one hour. (Fig. 4A). At 20 hps, the time of autonomously spawning under constant light conditions, no eggs had yet been spawned, and 23% of jellyfish spawned within the hour following exposure to light. At 24 hps, 20% of jellyfish maintained in the dark had spawned, with a further 39% spawning within the following hour. By 28 hps, most jellyfish (74%) kept in darkness had completed spawning, with the remainder spawning within the hour (Fig. 4B). In these experiments, although most oocytes were released within one hour from the first ovulation event, egg release continued for at least three hours (Fig. 4C). Overall, these observations show that ovulation is asynchronous and takes longer when *Clytia* sp. IZ-D jellyfish are maintained in a dark environment compared to those under light-dark cycles or constant light. This suggests that light plays a second role in maintaining a synchronous circadian spawning cycle, in addition to modulating ovulation timing through light stimulus.

**Figure 4.**
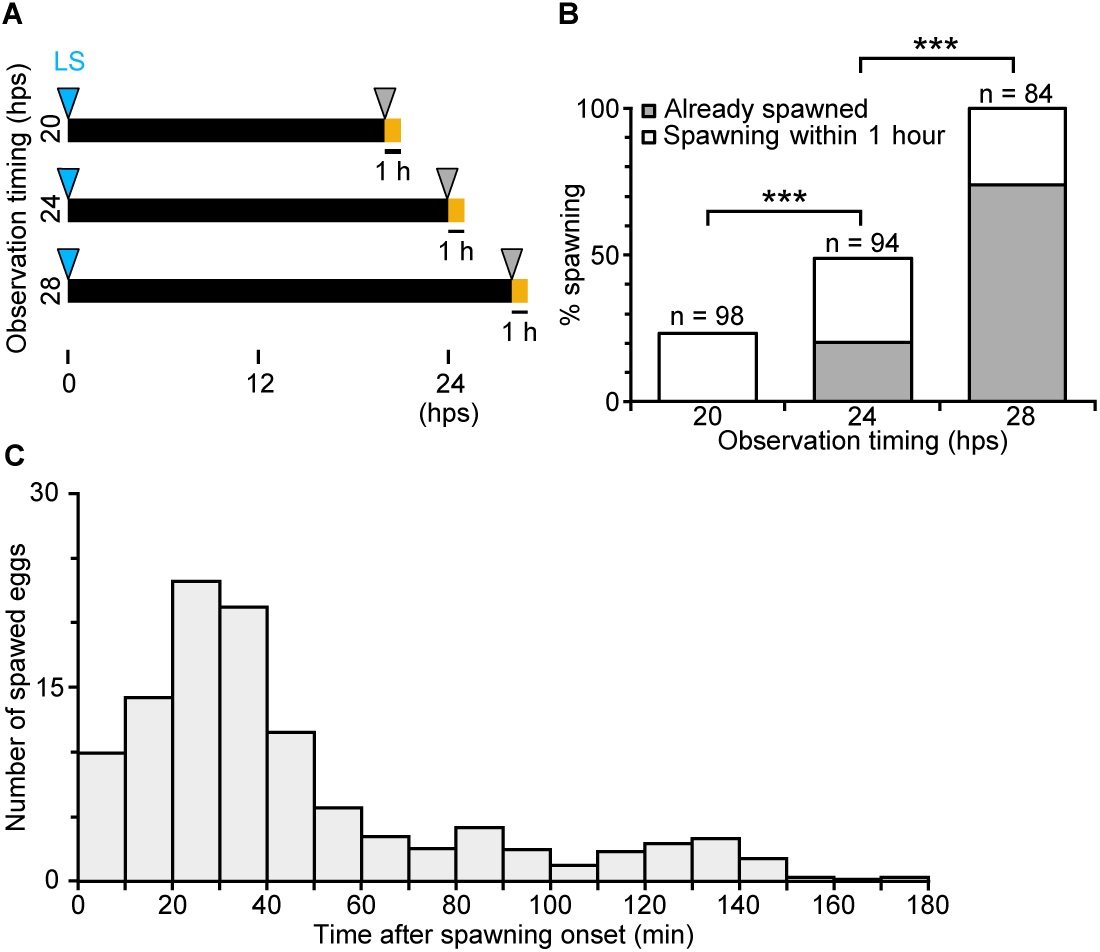
Asynchronous and spontaneous egg spawning from jellyfish under constant dark. (A) Experimental design to count spawning from jellyfish maintained in complete dark after the last spawning (LS). Jellyfish maintained in the dark were exposed to light at 20, 24 or 28 hps, and spawning before or one hour after light exposure was scored. (B) Percentage of jellyfish that had already started to spawn prior to light exposure (grey box) and those that started one hour after light exposure (white box). *n* = number of jellyfish. (C) Individual eggs released from jellyfish (N = 10) maintained in constant dark, counted every 10 minutes after the first egg release from each jellyfish.

### A threshold MIH concentration dictates the timing of maturation

We speculated that their quasi-circadian spawning could be achieved in a circadian clock-independent manner through minor modifications of the opsin-induced MIH release and response mechanisms described previously in *C. hemisphaerica* (Quiroga-Artigas et al., 2020, 2018). Under this hypothesis, spawning time could depend on two limiting factors: the accumulation of released MIH in the gonad and the competence of individual oocytes (or inactive sperm) to respond to it. Spawning would occur when both factors exceed critical thresholds, which in *Clytia* sp. IZ-D jellyfish would be at around 22 hps, resulting in spawning at around 24 hps. To explore this hypothesis, we first determined the effective dose of MIH required to trigger maturation. Fully grown oocytes (stage III; Suppl. Fig. 2) were isolated from jellyfish cultured in the standard L12/D12 Zeitgeber at 22 hps (12 hours after the light stimulus, just before the anticipated timing of MIH-triggered oocyte maturation) and tested their response to synthetic WPRPamide, one of the MIH peptides functional in both *C. hemisphaerica* and *Cladonema pacificum* (Fig. 5A). The timing of events was compared with that of oocytes within the intact ovary, where GVBD (Germinal Vesicle Breakdown, indicating entry into first meiotic M phase) occurred between 22 to 22.5 hps, and second polar body formation (IIPBF), indicating completion of meiosis, within 90 minutes (Fig. 5B). WPRPamide at concentrations of 10^-8^ M or higher consistently triggered oocyte maturation in isolated oocytes, whereas at 10^-9^ M only a very low proportion of treated oocytes matured (Fig. 5C). These results imply that oocytes receive an MIH signal after about 22 hps to initiate the maturation and spawning process.

**Figure 5.**
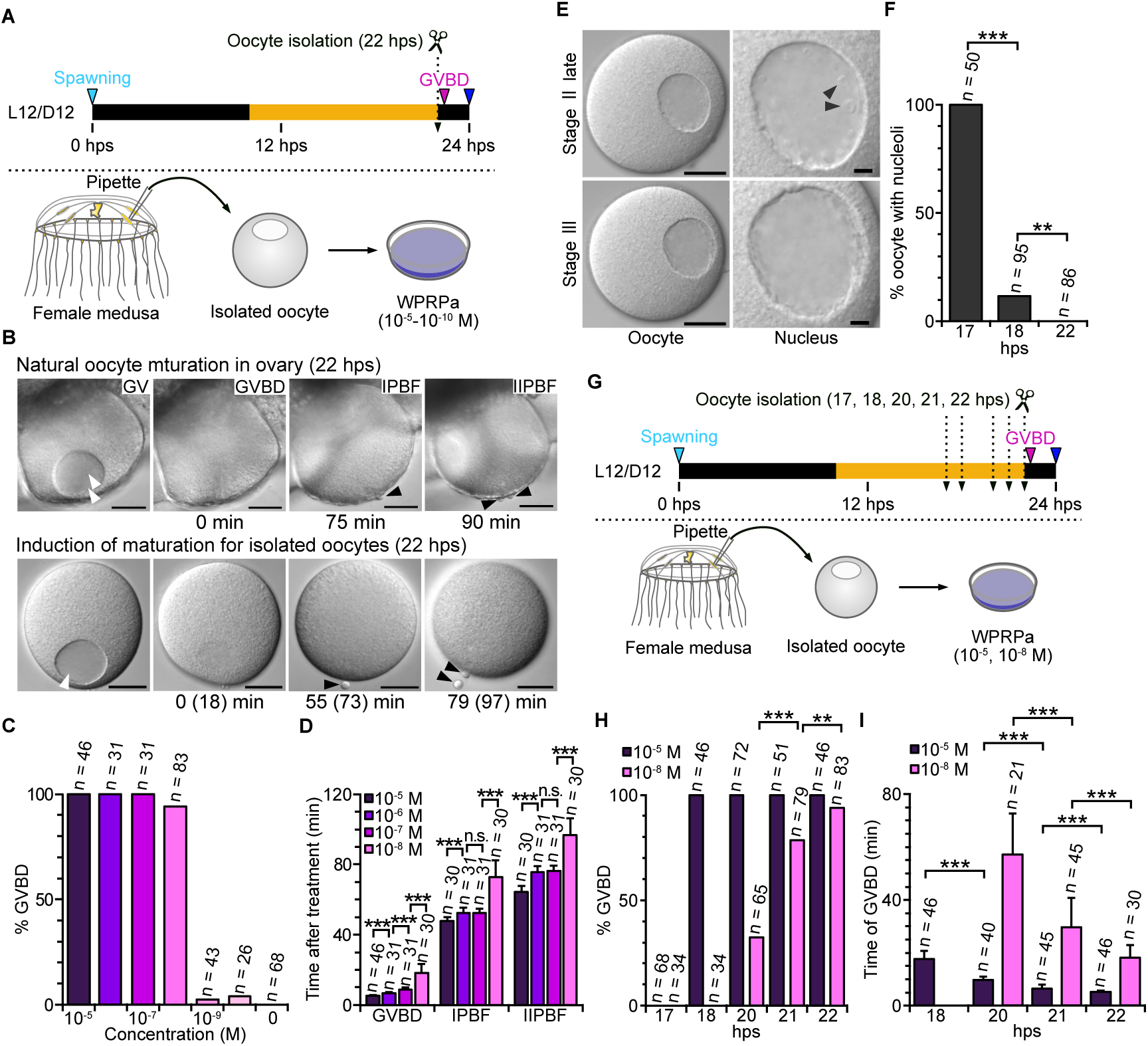
Oocyte development and acquisition of MRC (MIH-reactive competence) (A) Experimental procedure; oocyte isolation at 22 hps, just before endogenous onset of oocyte maturation (meiotic resumption), followed by in vitro induction of maturation with WPRPamide, employed in (B–D). (B). Stages of oocyte maturation in the gonad of jellyfish under L12/D12 cycle and in vitro treated with WPRPamide (10^-8^ M); GV: Stage III oocyte with germinal vesicle. GVBD: germinal vesicle breakdown; IPBF/IIPBF: first and second polar body formation. Times from GVBD are indicated below each panel. Numbers in parentheses show the time counted from WPRPamide addition. (C). Percentage of isolated oocytes that underwent GVBD upon WPRPamide treatment. (D) Time from WPRPamide addition to GVBD, IPBF and IIPBF. (E) Characteristic morphologies of late stage II oocytes (17 hps) and stage III (18 hps), with a higher magnification of the GV on the right. Stage III is defined by the absence of nucleoli, labelled by triangles. See also Supplementary Figure 2 for oocyte stages and their development over time. (F) The proportion of grown oocytes with nucleoli, which are indicative of stage II. Absence of nucleoli is characteristic to stage III. (G) The experimental procedure of oocyte isolation at different stages (17–22 hps) and induction of maturation by WPRPamide treatments used in (H) and (I). (H) Percentage of oocytes that underwent oocyte maturation in response to high (10^-5^) and low (10^-8^ M; likely equivalent to the endogenous MIH signal) WPRPamide, after isolation at different stages. (I) Time required from adding WPRPamide to seawater to GVBD. Time required for GVBD following 10^-8^ M WPRPamide treatment progressively decreased between 20 and 22 hps.

Two further observations support the conclusion that the effective MIH concentration acting in vivo is equivalent to 10^-8^ M WPRPamide applied in external seawater. Firstly, the sensitivity of *Clytia* sp. IZ-D oocytes to WPRPamide was similar to that of *C. hemisphaerica* and *Cladonema pacificum*, where WPRPamide induced oocyte maturation at 10^-7^ M but not at 10^-9^ M or less (Takeda et al., 2018). Secondly, the time required to complete oocyte maturation decreased when increasing WPRPamide concentrations (Fig. 5D). Notably, the duration from GVBD to IPBF and from IPBF to IIPBF shortened with higher peptide concentrations (Suppl. Video 3). Thus, the average duration from GVBD to IIPBF following treatment with 10^-8^ M WPRPamide (80 minutes) was closer to that observed during light-induced oocyte maturation in the gonad (90 ± 10 minutes) than to that induced by 10^-5^ to 10^-7^ M WPRPamide (60 to 70 minutes).

### Oocyte competence to respond to MIH increases progressively from 18 hps

We next examined the progression of oocytes through the final stages of growth and their competence to respond to WPRPamide when isolated at successive times between 17 to 22 hps. *Clytia* female gonads contain oocytes at different stages, with populations of vitellogenic oocytes progressing through growth stages I to III each day (Amiel and Houliston, 2009; Munro et al., 2023). We focused on late stage II and stage III (‘fully grown’) oocytes, both characterised by the peripheral positioning of the oocyte nucleus (termed GV for germinal vesicle), but distinguishable by the presence or absence of nucleoli (Suppl. Fig. 1). In *Clytia* sp. IZ-D female jellyfish maintained at 21°C, late stage Il oocytes were first detected at 17 hps, and stage III oocytes at 18 hps (Suppl. Fig. 1). At 17 hps, all oocytes exhibited nucleoli, indicating late stage II, while by 18 hps, most of the oocytes had lost their nucleoli, reaching stage III (Fig. 5F). To compare the competence of stage II versus stage III oocytes to respond to MIH, we isolated oocytes at successive times and treated them with WPRPamide (Fig. 5G, H). Oocytes isolated at 17 hps (i.e., all at late stage II) failed to respond to WPRPamide at any concentration. Oocytes isolated at 18 hps (Suppl. Fig. 2) underwent GVBD following treatment with high concentrations (10^-5^ M) of WPRPamide but did not respond to 10^-8^ M (Fig. 5H). Oocytes gradually acquired responsiveness from 20 hps onwards, such that virtually all oocytes responded to 10^-8^ M MIH at 21– 22 hps, just prior to the onset of maturation under standard L12/D12 conditions. The speed of maturation (measured as time from MIH treatment to GVBD) progressively decreased with oocyte age for a given WPRPamide concentration (Fig. 5I). Taken together, these results from MIH treatments of isolated oocytes allow us to define the capacity of stage III oocytes to respond to 10^-8^ M WPRPamide, which mimics the effective endogenous MIH concentration (see above), as MIH-Reactive Competence (MRC). The timing of MRC acquisition (i.e., 22 hps) coincides with the endogenous onset of oocyte maturation (22–22.5 hps).

### MIH-reactive competence develops autonomously with oocytes from 17 to 22 hps

To test whether oocytes require the gonadal environment to acquire MRC, we isolated late stage II oocytes at 17 hps from jellyfish maintained under a standard L12/D12 cycle, and subsequently cultured them in seawater (Fig. 6A). Isolated oocytes lost nucleoli, indicating progression from late stage II to stage III, within 45 minutes after isolation (Fig. 6B). As described above, late stage II oocytes immediately after dissection could not initially respond to MIH (Fig. 5H). After one hour of incubation in seawater, they become reactive to 10^-5^ M WPRPamide (Fig. 6C) but not to 10^-8^ M. After four to five hours of incubation, they become reactive to 10^-8^ M WPRPamide, with GVBD occurring with the endogenous timing (Fig. 6C, D), and thus can be considered to have acquired MRC. This timing corresponds to 21 and 22 hps. We conclude that MRC acquisition of isolated oocytes is not significantly different between oocytes isolated at 17 hps and 22 hps. This indicates that the oocytes from 17 hps have been programmed to acquire MRC autonomously at 21–22 hps.

**Figure 6.**
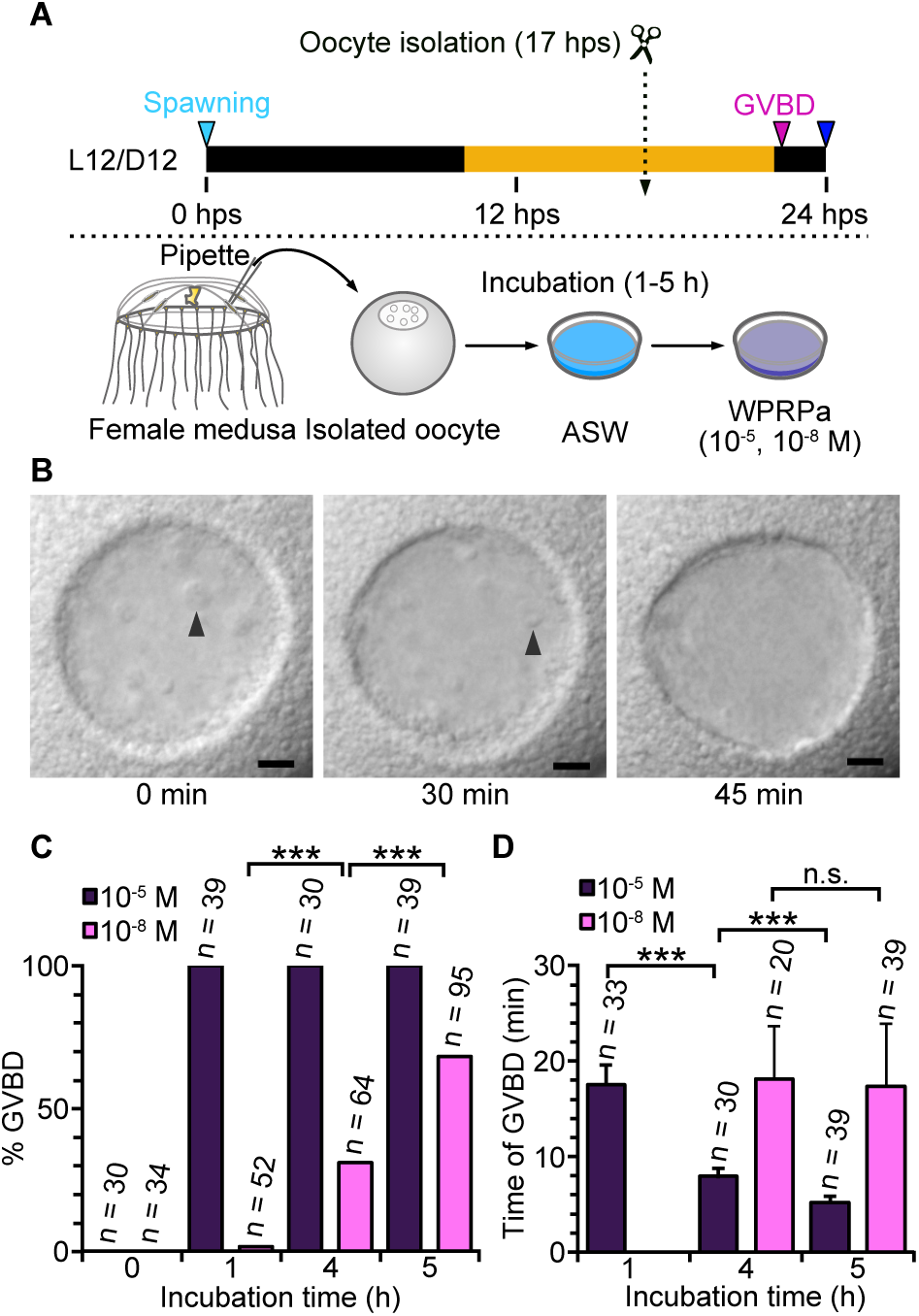
Late stage II oocytes autonomously develop to stage III and acquire MRC in vitro. (A) The experimental procedure. Late stage II oocytes isolated at 17 hps from the gonad were incubated in seawater for up to 5 hours (corresponding to 22 hps), then treated with WPRPamide. (B) GV morphology during culture. Arrowheads indicate nucleoli, which disappeared within 45 minutes. (C) Percentages of oocytes underwent oocyte maturation with WPRPamide, before (0 hour) and after (1– 5 hours) in vitro culture. (D) Time from WPRPamide addition to GVBD.

### Oocyte development and MRC acquisition are differentially affected by light

To distinguish the respective roles of MRC acquisition and oocyte development in the timing of oocyte maturation and spawning, we tested the MRC of oocytes isolated from jellyfish cultured under constant light (Fig. 7A–D) and constant dark (Fig. 7E–H). Under these conditions, natural ovulation occurs at 20 hps under constant light and asynchronously between 20 to 28 hps under constant darkness. Oocytes from jellyfish cultured under constant light conditions at 17 hps mostly exhibited nucleoli (i.e. were at late stage II), while by 18 hps the majority had lost nucleoli (i.e. reached stage III) (Table 1, Fig. 7B). The timing of the transition from late stage II to stage III was thus equivalent to that observed under L12/D12 conditions (Fig. 5E, F). In contrast to the L12/D12 Zeitgeber results, however, almost all oocytes isolated as early as 18 hps from constant-light jellyfish responded to both 10^-5^ and 10^-8^ M WPRPamide (100% and 92% respectively). Thus, under these conditions, most oocytes acquire MRC immediately after reaching stage III. This timing of MRC acquisition assayed in isolated oocytes coincides with the onset of oocyte maturation in intact jellyfish (18 hps; ovulation at 20 hps). This observation confirms that MRC acquisition can be uncoupled from morphological transition to stage III and, importantly, rules out the possibility that it is merely a function of growth time since the last spawning. Instead, it suggests that MRC acquisition is influenced by factors from the gonad under the control of the light-dark Zeitgeber.

**Figure 7.**
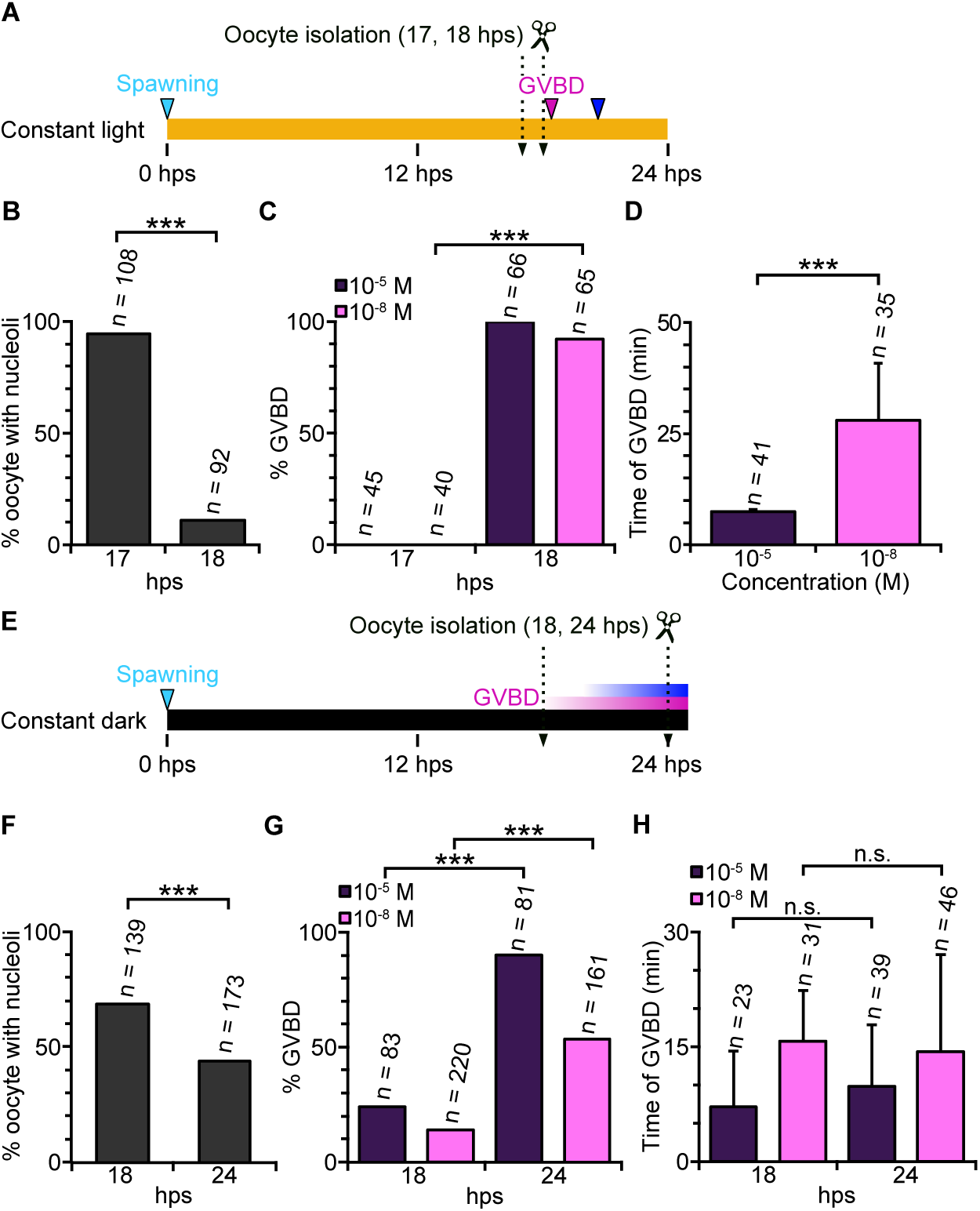
Timing of MRC acquisition of isolated oocytes mirrors that of spawning. (A–D) Experiments under constant light. (A) Timing of oocyte isolation; GVBD and spawning (18–18.5 hps and 20 hps, respectively) are indicated by magenta and blue triangles. (B) Percentage of isolated oocytes containing nucleoli, indicative of stage II. (C) Percentage of oocytes that underwent GVBD following WPRPamide treatment. (D) Time from WPRPamide addition to GVBD for oocytes isolated at 18 hps. (E–H) Experiments under constant dark. (E) Timing of oocyte isolation. The timings of asynchronous GVBD and spawning events are indicated by magenta and blue bars. (F) Percentage of isolated oocytes with nucleoli. (G) Percentage of oocytes that underwent GVBD following WPRPamide treatment. (H) Time from WPRPamide addition to GVBD for oocytes isolated in panel G.

Under constant-dark conditions, unlike under constant light, morphologically defined oocyte development was disrupted. Many of the grown oocytes isolated at 18 hps (32%) had lost their nucleoli, while at 24 hps, 44% were still at late stage II (Fig. 7F). Despite this asynchrony, the same relationship between MRC acquisition and spawning was observed as under the other light conditions, such that the proportion of oocytes responsive to 10^-8^ M WPRPamide also increased progressively (Fig. 7G). These observations using isolated oocytes mirror the asynchronous and long-lasting egg release observed from intact jellyfish under constant dark (Fig. 4). The detection of MRC in approximately half of stage III oocytes isolated from dark-maintained jellyfish as early as 18 hps (14% with MRC and 32% at stage III; Fig. 7G), albeit in lower proportions than for oocytes from constant-light conditions (92% MRC and 89% stage III; Fig. 7C), contrasts with the absence of MRC until 20 hps under L12/D12 Zeitgeber (at 18 hps: 0% MRC and 88% stage III; Fig. 5H). Thus arrival at stage III and MRC can be uncoupled.

Together, these observations suggest that attainment of stage III and acquisition of MRC are controlled by light in two distinct mechanisms. On one hand, light exposure prior to 17 hps favours the synchronous development of oocytes to reach stage III, characterised by the loss of nucleoli. On the other hand, the timing of MRC acquisition after reaching stage III is influenced by the timing of the previous dark-light cue experienced by the jellyfish, which can delay spawning by up to four hours (Fig. 8).

**Figure 8.**
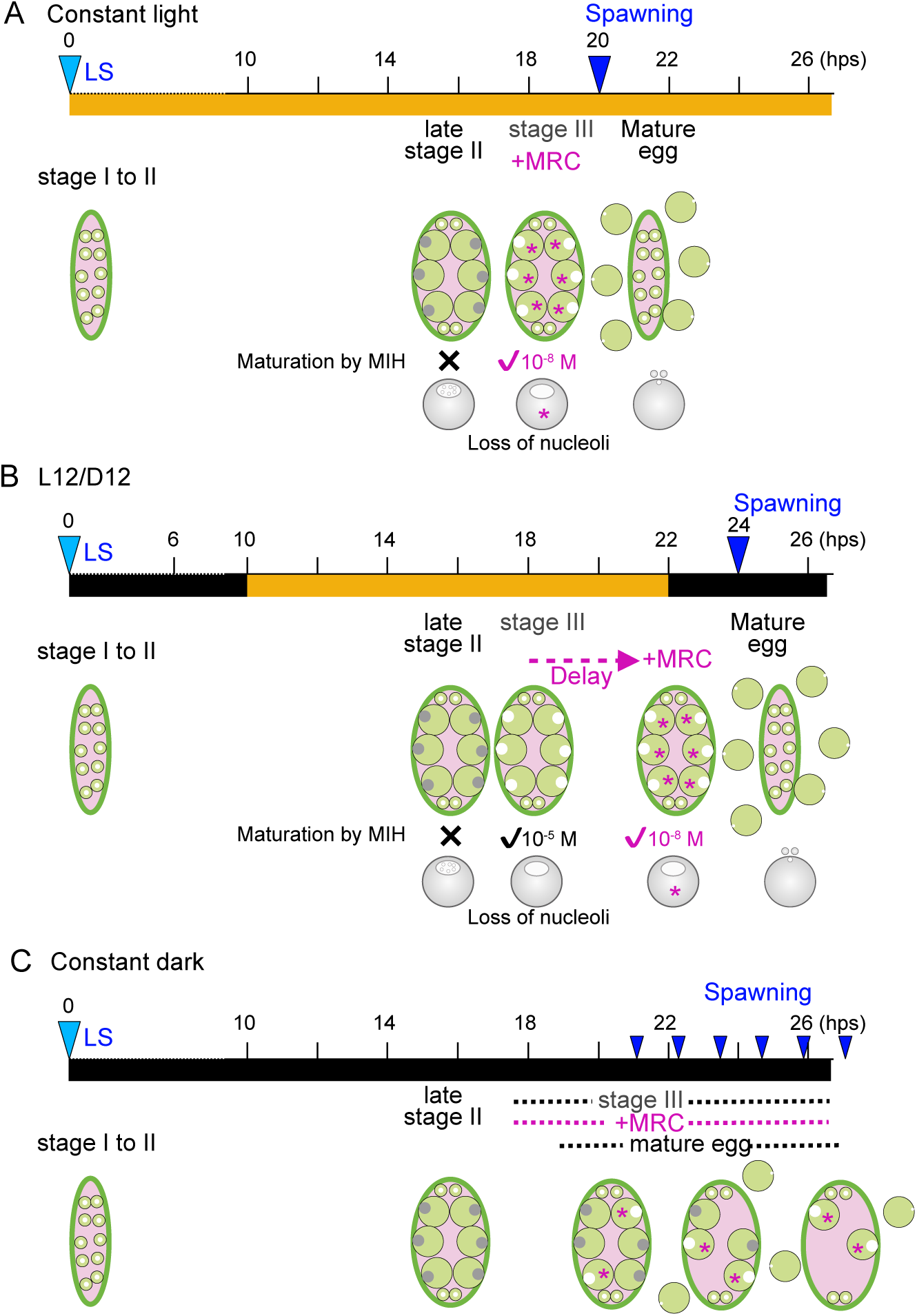
Comparison of the timing of oocyte development, MRC and spawning under three light/dark regimes. (A) In constant light, synchronous ovulation occurs every 20 hours. Both stage III arrival and MRC occur at 18 hps, about two hours before the spawning. (B) Under a L12/D12 cycle, synchronous spawning occurs every 24 hours, 14 hours after the light stimulus. Stage III arrival and MRC occur at 18 and 21–22 hps, respectively, with synchronous spawning at 24 hps. (C). In constant darkness, spawning occurs but is asynchronous. MRC: MIH-reactive competence, defined as the ability to respond to 10^-8^ M exogenous WPRPamide, deduced to match the endogenous MIH level to trigger spawning. Previous spawning (LS) is indicated with light blue triangles. Synchronous (A and B), and asynchronous (C) spawning events are shown with dark blue triangles.

## Discussion

From the Pacific coast of Japan, we identified a new species of hydrozoan jellyfish *Clytia* sp. IZ-D, which releases gametes at dusk in the natural habitat. In this study, by observing egg release from jellyfish, we revealed that *Clytia* sp. IZ-D possesses an autonomous circadian rhythm of synchronous gamete release every 20 hours at 21°C under constant light conditions, as summarised in Fig. 8A. The periodic cycle length is temperature-dependent. Under light-dark cycle, synchronous ovulation becomes adjusted to a 24-hour period, in which gamete release occurs synchronously 14 hours after the preceding dark-to-light transition at 21°C (Fig. 8A), requiring a minimum of four hours of dark followed by light illumination. The regulation of gamete release by this dark-to-light cue is not fully determinative but is constrained within a range of 20 to 24 hours from the previous spawning event. In females, the lower limit is defined by a minimal period of 18 hours required for development for the daily cohort of growing oocytes. Thus, in this species, the role of light stimulus can be considered as modulating an autonomous circadian ovulation cycle. This is the first observation that hydrozoan jellyfish display an autonomous circadian rhythm controlling precise and synchronised gamete release. Under constant darkness, autonomous egg maturation and release occur asynchronously, starting from 20 hours after the previous ovulation. Light therefore plays an additional, permissive role in maintaining the autonomous circadian rhythm.

Our observations show that key determinants of the autonomous ovulation cycle are oocyte growth and the acquisition of maturation competence. The shortest ovulation interval (20 hours at 21°C) is constrained by the time required for growing oocytes to develop to stage III, the stage at which they are ready to undergo meiotic maturation, occurring at 18 hps. The actual timing of ovulation correlates with the timing of the oocytes to acquire the competence to respond to 10^-8^ M WPRPamide in the external seawater (MRC). This concentration simulates the effective endogenous MIH in the gonad, based on the estimation derived from the duration between GVBD and PBFI/PBFII. Assuming this level is present in the gonad from at least 18 hps (see below), oocyte growth and acquisition of MRC would be sufficient to determine the next oocyte maturation and ovulation schedule and thus to account for the autonomous circadian gamete release observed in constant light in *Clytia* sp. IZ-D.

### MIH-induced oocyte maturation is conserved

MIH-induced oocyte maturation has been demonstrated in a number of hydrozoan species, including *Clytia hemisphaerica*, which is closely related to *Clytia* sp. IZ-D, and *Cladonema pacificum*, both of which utilise nearly identical peptide sequences, respectively RPR(P/A)amide and (W/R)PRPamide, as the endogenous MIH (Takeda et al., 2018). In *C. hemisphaerica*, neurosecretory cells in the gonad ectoderm expressing an opsin, CheOpsin9, release MIH upon light stimuli. Oocytes receive MIH and undergo maturation (completion of meiosis), beginning with germinal vesicle breakdown (GVBD) 20–25 minutes after light stimulation or MIH addition to isolated oocytes. This process completes within 90–120 minutes (Amiel and Houliston, 2009). The timeline from GVBD to the completion of meiosis is similar in *Clytia* sp. IZ-D suggesting that MIH signalling in this species is also received by oocytes approximately two hours before ovulation.

We do not yet know when and how MIH is released into the gonadal extracellular space in *Clytia* sp. IZ-D. It does not seem likely that a pulse of MIH is released simply by counting a delay of exactly 12 hours after light stimulation or 22 hours from the previous ovulation. BMAL1/CLOCK circadian clock genes were likely lost in a common hydrozoan ancestor (Kanaya et al., 2019), so this mechanism is not available to control such precise timing. It also appears highly improbable, although it still needs to be examined, that a novel circadian clock mechanism equivalent to the BMAL1/CLOCK system arose within Hydrozoa and is employed in *Clytia* sp. IZ-D to induce oocyte maturation. We propose that the characteristic spawning cycle of *Clytia* sp. IZ-D instead involves gonad neurosecretory cells expressing opsins and mediating MIH release, as in *C. hemisphaerica,* but operating with distinct kinetics. One possibility is that MIH is slowly released into the extracellular space during light conditions from MIH-expressing neurosecretory cells, and that oocytes only respond once they have activated sufficient G-protein-coupled MIH receptors (MIH-R (Quiroga-Artigas et al., 2020)) on the oocyte surface to elicit the downstream response. However, slow MIH release alone is unlikely to explain the precision of ovulation timing, as it would require implausibly precise control to reach a receptor-binding threshold with such accuracy in the absence of a timer. Our current hypothesis is that the crucial elements determining spawning timing are oocyte growth and competence acquisition to respond to MIH (Fig. 8), which could be released slowly and/or constitutively in the presence of light.

### Oocyte growth as a timer component

As outlined above, our observations suggest that the competence of oocytes to respond to MIH serves as a key control point in defining the timing of oocyte maturation. We identified two light-dependent processes: the transition of oocytes to stage III and the acquisition of MRC. The arrival at stage III is characterised by the loss of nucleoli and readiness to undergo meiotic maturation, likely coinciding with the partial chromosome condensation observed at diakinesis stage (Munro et al., 2023). This occurs at 18 hours after the previous spawning (hps) in all conditions except for continuous dark (Fig. 8). This partly explains why ovulation does not occur at 20 hps or earlier, and indicates that light exposure during the growth period is necessary for synchronous oocyte development to stage III. Such a minimum delay for spawning to allow oocyte growth has also been observed in other hydrozoan species. Future studies in other species will test whether exposure to light favouring synchronous oocyte development is a common feature in hydrozoans.

Another control point, the acquisition of MRC, is influenced by light and, more specifically, by the timing of a preceding dark–light transition. The response of oocytes isolated from gonads to MIH suggests that the timing of MRC acquisition, which varies from 18 to 22 hps, is already autonomously programmed in the oocytes by 17 hps, depending on the timing of the dark–light stimulus before this stage. One possible model that can explain the 14-hour delay from the onset of the light period to oocyte maturation is that oocytes acquire MRC progressively over time, and that a transition to light following four or more hours of darkness stalls this process for several hours (Fig. 8B). This MRC acquisition process appears to be controlled independently from the transition from late stage II to III, which requires an exposure to light (Fig. 8), but is not affected by the dark–light transition. This may explain why ovulation timing varies among oocytes in the same gonad, up to 40 to 60 minutes under L12/D12 Zeitgeber conditions and constant light (Fig. 2H and 3B). Once MIH activity in the gonad has reached the equivalent 10^-8^ M of WPRPamide applied externally, individual oocytes will react to it as soon as they acquire MRC.

It will be fascinating to address the molecular basis of the light modulation of gamete development and MRC in future studies. One speculation is that the gradual increase of MRC in oocytes may be stimulated by continuous weak signalling from light-sensitive neurosecretory cells in the gonad ectoderm, contrasting with the immediate release of MIH triggered by CheOpsin9 activation to stimulate immediate oocyte maturation upon light stimulus in *C. hemisphaerica*. The molecular nature of such a continuous signal could potentially be mediated by MIH itself and/or other neuropeptides. One possibility is that the cAMP signalling downstream of MIH/MIH-R, which induces oocyte maturation, also promotes oocyte development. Consistent with this hypothesis, treatment of in vitro isolated late stage II oocytes of another hydrozoan, *Cytaeis uchidae,* with the membrane-permeable cAMP analogue Br-cAMP at a concentration lower than necessary to initiate oocyte maturation, triggers the loss of nucleoli that marks the transition to stage III (Takeda, unpublished observation). In this gradual MIH-accumulation hypothesis, light-induced delay of MRC acquisition (e.g. delay of MRC/spawning under L12/D12 Zeitgeber compared to continuous light) may be explained by attenuation of the slow MIH release several hours after light stimuli, or by the release of antagonistic neuropeptides upon light stimulus. In parallel, considering that light stimulus delays spawning only when applied between 6 and 10 hps, it will be particularly interesting to investigate which oocyte development process during prophase I arrest is modulated by the light stimulus.

A similar mechanism is likely employed for sperm release regulation. Sperm release timing occurs slightly earlier (roughly 13 hours after the light stimulus) than female ovulation. However, little is known about how sperm release is controlled. In *C. hemisphaerica*, unlike fully grown oocytes, which are arrested at the end of prophase I, spermatozoa that have completed meiosis accumulate in male gonads (Munro et al., 2023), before being activated by MIH to become motile and be released from the gonad (unpublished observation). In the coral *Astrangia poculata,* sperm activation is induced by cAMP signalling (Glass et al., 2023), which is likely the second messenger of the seven transmembrane MIH receptor MIH-R, expressed during spermatogenesis as well as oogenesis in *Clytia* (Quiroga-Artigas et al., 2020)

### Evolutionary and ecological advantages

For marine species that externally release and broadcast gametes, timing control is crucial to successfully achieving sexual reproduction. In both *C. hemisphaerica* and *Clytia* sp. IZ-D, gamete release is coordinated, with male sperm release starting slightly earlier than female ovulation (Munro et al., 2023). Sperm release tends to continue longer than ovulation in both *Clytia* species (Kitsui, unpublished observation). Such a prolonged sperm broadcast strategy may be an adaptation to sperm competition (Lotterhos and Levitan, 2010) among jellyfish sparsely distributed in the water.

In *C. hemisphaerica*, fertilisation success requires gamete mixing within one hour after egg release. Sunrise and sunset are the few environmental cues that allow animals at a distance to synchronise gamete release. Most known hydrozoan jellyfish release gametes within a few hours of light stimulation (Freeman, 1987; Freeman and Ridgway, 1988; Roosen-Runge, 1962; Takeda et al., 2006). Multiple corals in the same regions will release gametes simultaneously, risking hybridisation (Willis et al., 2006). Changing spawning times may confer an advantage by avoiding cross-species fertilisation, and a genetic change that alters the timing of gamete release within a population could reproductively isolate it from the others. This, in turn, could lead to the eventual establishment of a new species (Westram et al., 2022). We identified at least three *Clytia* species, including *Clytia* sp. IZ-D in the bay of Izuashima Island (Kitsui and Deguchi, unpublished observations). The adaptation of gamete release timing to dusk, potentially through the innovation of an alternative circadian clock, might have allowed the speciation of *Clytia* sp IZ-D from the common ancestor shared with other *Clytia* species. Another potential ecological advantage of an autonomous circadian cycle is the robustness it confers against seasonal climate instability. *Clytia* sp. IZ-D jellyfish can continue synchronised gamete release even if the sunlight does not follow a regular 24-hour cycle, for example, during the seasonal typhoons in summer and autumn – common in this region – or under the influence of artificial light sources.

## Materials and Methods

### Collection and cultures of biological materials

Medusae of the genus *Clytia* (*Clytia* sp. IZ-D, *Clytia* sp. IZ-C) were collected through a series of regular sampling in Izushima Island (Miyagi Prefecture, Japan) from July to October 2023. Collection was performed using the jellyfish net described previously (Deguchi et al., 2005), consisting of a 1 mm nylon mesh filter attached to a 300 mm diameter metal ring frame. The net was cast from the shore and slowly dragged at a constant speed to target plankton located less than 2 m below the sea surface, with sampling repeated over several hours. Collected jellyfish were transferred into a stainless steel cooking container with seawater and identified morphologically.

### Local climate data

Average monthly water temperature data from Enoshima Island station (8 km southwest of Izushima Island) were provided by the Miyagi Prefecture Fisheries Technology Institute (https://tohokubuoynet.myg.affrc.go.jp/Vdata/). Sunlight data for Sendai city (65 km west of Izushima) for 2024 was obtained from the Ephemeris Computation Office of the National Astronomical Observatory of Japan (https://eco.mtk.nao.ac.jp/koyomi/dni/dni04.html.en).

### Phylogenetic analysis of 16S rRNA sequences

DNA was extracted from medusae using the NucleoSpin Tissue XS (MACHEREY-NAGEL) according to the manufacturer’s protocol and used as a PCR template. PCR was conducted using KOD One PCR Master Mix (TOYOBO), with PCR primers 16S-F1 (5’-ACGGAATGAACTCAAATCATGTAAG-3’) and 16S-R1 (5’-CCTTTTGTATAATGGATTTACAAG-3’). PCR cycle was as follows: initial denaturation at 98°C for 1 minute, followed by 35 cycles of 10 seconds at 98°C, 5 seconds at 50°C or 52°C and 5 seconds at 68°C, with a final extension at 68 °C for 1 minute. The PCR products were purified using Nucleospin Gel and PCR Clean-up kit (MACHEREY-NAGEL) and sequenced at Eurofins Genomics (Tokyo, Japan). The sequences obtained were aligned using MUSCLE in the MEGA 11 software for macOS (Stecher et al., 2020; Tamura et al., 2021). The alignment data of 16S rRNA, including their accession numbers, is provided in Supplementary File 1 in FASTA format. Phylogenetic trees were generated using the maximum-likelihood (ML) method with 1000 bootstrap replicates, based on the K2P model. The sequence of *Cladonema pacificum* (AB720901.1) for 16S rRNA was used as the outgroup. Bootstrap values greater than 50% are shown above the branches as node support values in Fig. 1C.

### Jellyfish culture and polyp colony establishment

Male and female jellyfish were maintained in polypropylene dishes (60 mm diameter, 35 mm height) with sealing lids, filled with artificial seawater (ASW: SEALIFE salt, 35 g/L 30 ppt, Marinetech, Japan). ASW was used for all in vivo experiments in this work. A pair of male and female jellyfish identified in October 2023 was placed together in a dish for crossing. The resulting planula larvae were transferred to a new dish, where they underwent natural metamorphosis to make polyp colonies. They were subsequently transplanted to individual dishes and placed in a larger tank with water flow maintained by fine bubble aeration. Jellyfish and polyp colonies were fed live *Artemia salina* nauplii once every day or two. Newly budded jellyfish were collected from the dishes and reared separately in other dishes. Under these culture conditions, medusae began releasing eggs and sperm within two weeks.

### Adjustment of light-dark cycle and observation of spawning timing

*Clytia sp.* IZ-D D-A1 and D-A3 strain jellyfish, less than three months after liberation from the polyp colonies, were used in this study. Mature jellyfish in culture dishes were placed in a 21°C incubator under a light-dark Zeitgeber cycle controlled by a programmable timer (Koizumi Computer or Ohm Electric Inc.) and illuminated with white LED light (Yazawa LE3WH, 0.2 W). Under standard culture/growth conditions, jellyfish were maintained on a 12-hour light and 12-hour dark cycle (L12/D12) at 21°C. After the last spawning event (marked in LS in the figures), each jellyfish was transferred to an individual well of a 6-well plastic dish (AsOne VTCP-6) and maintained in another incubator (Mitsubishi Electric CN-40A) shifted to the experimental Zeitgeber. The timing of egg and sperm release for each jellyfish was defined by the initiation of release and measured through continuous observation using a stereomicroscope (Leica M165C) or an inverted microscope (Nikon Eclipse TE300). In the case of spawning during the dark period, where observation under a light microscope might disturb spawning, we first estimated the approximate timing of spawning by pilot experiments conducted without light. Precise observations were then made under light microscopes starting 40 minutes before the anticipated spawning time, at which point oocyte maturation had already commenced, thus minimising any impact on oocyte maturation timing. We also confirmed that the interruption of dark periods by this observation did not affect the ovulation timing on the following day. For jellyfish maintained under constant dark conditions, where spawning timing was highly variable among jellyfish and oocytes, and where the light exposure would interrupt the dark environment, jellyfish were sampled at different time points, and the spawning by the time of exposure and in the following hour was observed. (See also the main text.)

### Oocyte isolation from the gonad and in vitro maturation

The jellyfish were anaesthetised in ASW containing Mg^2+^ (a 1:1 mix of 0.53 M MgCl_2_ and ASW) during manipulation. Oocytes were isolated from the gonad of female jellyfish under an upright microscope (Nikon Eclipse 80i) by aspirating with a microcapillary pipette, prepared by pulling a Pasteur pipette under a Bunsen burner flame and cutting the tip to an opening of 0.2 mm. Immediately after isolation, oocytes were staged based on nuclear morphology and the presence and number of nucleoli (see Suppl. Fig. 2 for staging criteria). Small oocytes (earlier than early stage II) were excluded. Isolated oocytes were maintained in plastic Petri dishes containing ASW. The peptide WPRP-NH_2_ (WPRPamide), the most potent isoform of the endogenous MIHs identified in *C. hemisphaerica* (Takeda et al., 2018) was synthesised by Genscript, dissolved into H_2_O at 10^-3^ M and stored at −20°C. Before use, it was diluted and added to the culture ASW at concentrations ranging from 10^−5^ to 10^−8^ M. Isolated oocytes were transferred to wells of a 24-well plate (Falcon 353047) containing the ASW supplemented with WPRPamide. The timing of oocyte maturation events, notably germinal vesicle breakdown (GVBD) and the formation of the first and second polar bodies (IPBF and IIPBF, respectively), was recorded through observation under the inverted microscope. Completion of oocyte maturation was defined by the formation of the second polar body. Experiments with isolated oocytes were performed in a room maintained at approximately 21°C.

## Abbreviations

MIH: Maturation-Inducing Hormone
hps: hours post-spawning
GVBD: Germinal Vesicle Breakdown
PBFI/PBFII: 1st and 2nd Polar Body Formation
MRC: MIH-Reactive Competence

