## Supplementary material for "A light-modulated clock mechanism in a hydrozoan jellyfish that synchronises evening gamete release": Suppl. File 1-3

Supplementary
Figure 1

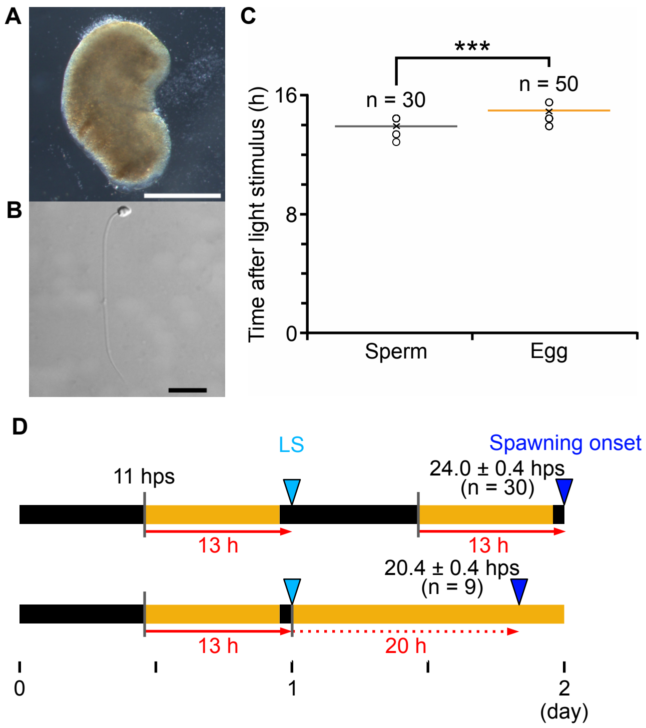


**Supplementary Figure 1. Sperm release from male jellyfish is coordinated with female ovulation.** (A) Isolated male gonads that release sperm. bar=0.5 mm (B) A high magnification image of a sperm. bar: 10 μm. (C) Comparison of male and female spawning time measured from the onset of light stimuli. Sperm release occurred earlier (13.0 ± 0.3 hours) than egg spawning (13.9 ± 0.3 hours). Sperm release timing is defined by its onset from a jellyfish, as it continues up to one hour. Bars indicate standard deviation *n*= number of jellyfish. (Welch's test, *** p ≤ 0.001) (D) Sperm release also exhibits regular spawning at dusk under a light cycle and autonomously under constant light.

Supplementary
Figure 2

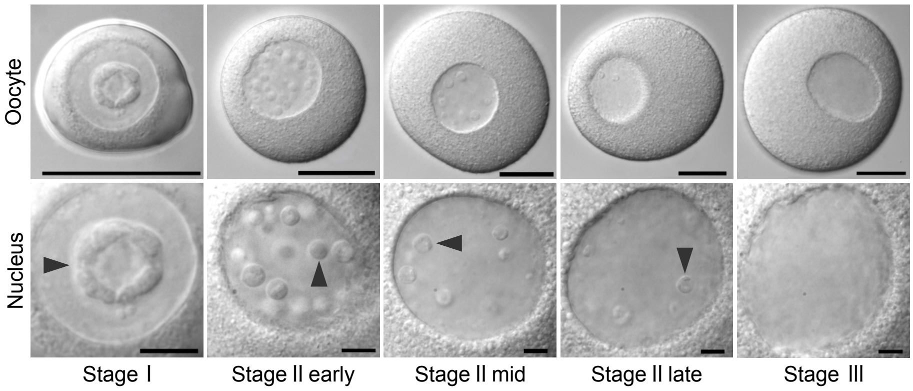


**B**

| Oocyte stage | Diameter (μm) | Nucleoli | | Response to WPRPa | Observable period (hps) |
|---|---|---|---|---|---|
| | | Number | Size | | |
| Stage Ⅰ | < 50 | Single | Large | No | 0-24 |
| Stage Ⅱ early | < 100 | > 30 | Small | No | 0-24 |
| Stage Ⅱ mid | < 150 | 10-20 | Small | No | 10-16 |
| Stage Ⅱ late | 160-180 | 3-10 | Small | No | 16-18 |
| Stage Ⅲ | 160-180 | No | | ○ | 18-22 |

**Supplementary Figure 2 Oocyte development.** (A) DIC images of oocytes and their nuclei at different stages. The staging criteria were based on Amiel et al., 2009 for *C. hemisphaerica*, with additional clarification that late stage II is defined by peripheral migration of GV and presence of nucleoli, while stage III is determined by the disappearance of nucleoli. Arrowheads indicate nucleoli. Bars are 50 μm for oocyte panels and 10 μm for nucleus panels. (B) Characteristics of oocyte stages and their occurrence after the previous spawning (hps).

Supplementary
Figure 3

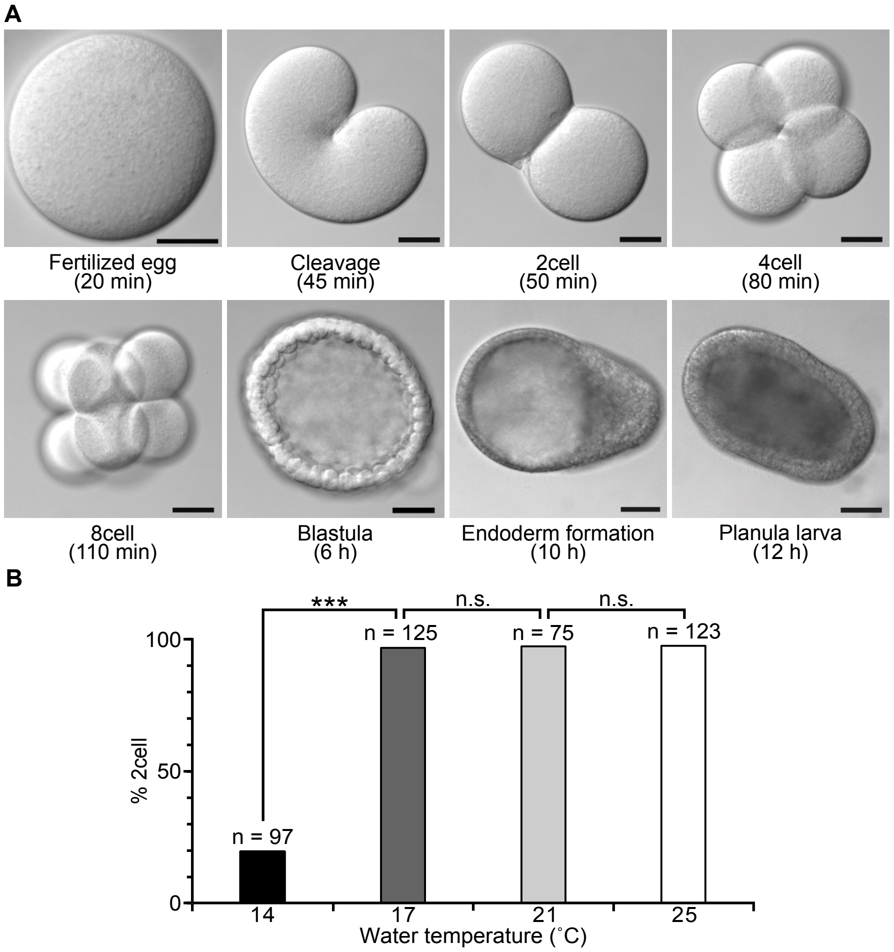


**Supplementary Figure 3. Development and fertilisation rates** (A) DIC images of each embryonic stage and planula larva. Scale bars: 50 μm. Time after fertilisation is indicated in parentheses. (B) Fertilization efficiency at different temperatures (chi-squared test, *** $p \leq 0.001$, n.s. $p \leq 0.05$)
